# Post-error slowing reflects the joint impact of adaptive and maladaptive processes during decision making

**DOI:** 10.1101/2021.12.22.473805

**Authors:** Fanny Fievez, Gerard Derosiere, Frederick Verbruggen, Julie Duque

**Affiliations:** Institute of Neuroscience, Université Catholique de Louvain, 1200, Brussels, Belgium; Department of Experimental psychology, Ghent University, 9000 Gent, Belgium

**Keywords:** speed-accuracy tradeoff, error processing, cognitive control, attention, emotion

## Abstract

Errors and their consequences are typically studied by investigating changes in decision speed and accuracy in trials that follow an error, commonly referred to as “post-error adjustments”. Many studies have reported that subjects slow down following an error, a phenomenon called “post-error slowing” (PES). However, the functional significance of PES is still a matter of debate as it is not always adaptive. That is, it is not always associated with a gain in performance and can even occur with a decline in accuracy. Here, we hypothesized that the nature of PES is influenced by one’s speed-accuracy tradeoff policy, which determines the overall level of choice accuracy in the task at hand. To test this hypothesis, we investigated post-error adjustments in subjects performing the same task while they were required to either emphasize speed (low accuracy) or cautiousness (high accuracy) in two distinct contexts (hasty and cautious contexts, respectively) experienced on separate days. Accordingly, our data indicate that post-error adjustments varied according to the context in which subjects performed the task, with PES being solely significant in the hasty context. In addition, we only observed a gain in performance after errors in a specific trial type, suggesting that post-error adjustments depend on a complex combination of processes that affect the speed of ensuing actions as well as the degree to which such PES comes with a gain in performance.

## 1) Introduction

We all make mistakes. For instance, many of us have experienced sending an email to the wrong person. After such an error, we typically write a second message to apologize and rectify. But when we send the second email, we usually take more time to check that the recipient is correct. We therefore adapt our behavior in order to avoid reproducing previous mistakes. Such an ability to adapt after an error is essential to achieve our goals.

Errors and their consequences are typically studied in two-choice reaction time tasks by investigating changes in decision speed and accuracy in trials that follow an error, commonly referred to as “post-error adjustments”. Using such tasks, many studies have reported that subjects slow down following an error, a phenomenon called “post-error slowing” (PES) (Fu et al. 2019, Dubravac et al. 2020, Nigbur and Ullsperger 2020, Topor et al. 2021).

The functional significance of PES is still a matter of debate though (Wessel 2018, Damaso et al. 2020, Kirschner et al. 2020). Because slowing down after an error is sometimes associated with an increase in accuracy, PES is traditionally attributed to adaptive adjustments of decision policies, favoring a more cautious response style to improve performance in the subsequent trial (Rabbitt and Vyas 1970, Smith and Brewer 1995, Cavanagh et al. 2014, Siegert et al. 2014, Purcell and Kiani 2016, Steinhauser and Andersen 2019, Beatty et al. 2020). However, several recent studies have revealed that PES can also occur in a somewhat ‘maladaptive’ way as slowing does not necessarily lead to improvement in accuracy; in fact, PES can even come with a decrease in decision accuracy (Ceccarini et al. 2019, Eben et al. 2020b, Kirschner et al. 2020, Schroder et al. 2020, Smith et al. 2020, Compton et al. 2021). These findings indicate that the functional significance of PES may vary according to the context in which it is observed.

A careful analysis of the literature reveals that the degree to which PES is adaptive (*i.e*., benefits accuracy) or ‘maladaptive’ (*i.e*., takes place without accuracy improvement) depends partly on the average level of accuracy of subjects in the task at play. That is, in studies reporting an adaptive PES, the overall level of choice accuracy is typically low (i.e. generally between 60 - 80% of correct choices) because the task is relatively complex and/or because the instruction requires subjects to respond quickly within a given time limit (Siegert et al. 2014, Purcell and Kiani 2016, Steinhauser and Andersen 2019). In this situation, errors are clearly expected and slowing down after them has a positive effect on choice accuracy (Hajcak et al. 2003, Siegert et al. 2014, Dyson et al. 2018, Wessel 2018, Damaso et al. 2020). By contrast, studies reporting a maladaptive PES rather use reaction time tasks that are quite simple such that the overall level of choice accuracy is usually much higher (i.e. more between 80 – 100% of correct choices) (Notebaert et al. 2009, Nunez Castellar et al. 2010, Houtman et al. 2012, Eben et al. 2020b, Kirschner et al. 2020, Li et al. 2020, Compton et al. 2021). In such settings, errors represent infrequent and unexpected events that may catch attention, resulting in a maladaptive PES that deteriorates (rather than enhances) choice accuracy in the consecutive trial (Sokolov 1963, Nunez Castellar et al. 2010, Houtman et al. 2012).

Thus, whether PES is adaptive or maladaptive might be partly influenced by choice accuracy. This in turn depends on task characteristics, such as the global task difficulty or the manipulation of the decision speed. Indeed, most decisions require balancing speed and accuracy, making the speed-accuracy tradeoff (SAT) a universal property of behavior (Henmon 1911, Rinberg et al. 2006, Salinas et al. 2014, Guo et al. 2019, Reynaud et al. 2020, Miletic et al. 2021). Humans and other non-human animals are able to adjust their SAT depending on context, favoring either hasty (i.e., high speed, low accuracy) or cautious (i.e., low speed, high accuracy) decision policies (Chittka et al. 2009, Heitz 2014, Spieser et al. 2017, Thura 2020). Hence, because choice accuracy varies depending on the SAT, it is plausible that PES can shift from being adaptive to being maladaptive depending on whether the emphasis is on speed or accuracy when performing the same task in separate blocks.

In conclusion, past research suggests that errors can trigger at least two types of PES, which are either adaptive or maladaptive (van Driel et al. 2012, Schiffler et al. 2017, Wessel 2018). These two different processes have been evidenced in separate studies using distinct tasks or instructions where performance is either characterized by a low or a high level of choice accuracy, respectively. Here, we hypothesized that the type of adjustments that follows an error can also vary within a given task depending on whether the SAT context favors a hasty (i.e., high speed, low accuracy) or a cautious (i.e., low speed, high accuracy) decision policy. More precisely, we predicted that errors would be common and expected when the context favors choice speed due to the choices’ promptness (Damaso et al. 2020), whereas they would be rare and unexpected when the context favors choice accuracy. Therefore, we expected adjustments to be less adaptive (and potentially maladaptive; i.e., slowing without an improvement in accuracy) when the emphasis is on choice accuracy in a cautious SAT context compared to when the emphasis is on response speed. To test this hypothesis, we used a modified version the “tokens task” (Cisek et al. 2009, Derosiere et al. 2019, Derosiere et al. 2021b), involving choices between left and right index fingers. In this task, incorrect choices led either to a *low* or to a *high* penalty in two different SAT contexts, inciting subjects to implement either *hasty* or *cautious* decision policies, respectively. We predicted that PES would be more adaptive in the low than in the high penalty context.

## 2) Material and Method

### Participants

A total of 43 healthy volunteers participated in this study (25 Women: 23.5 ± 2.3 years old). All participants were right-handed according to the Edinburgh Questionnaire (Oldfield 1971). None of them had any neurological disorder or history of psychiatric illness or drug or alcohol abuse; and no one was following any clinical treatment that could have influenced performance. Participants were financially compensated for their participation and could also receive extra compensation based on their performance on the task (see below). All gave written informed consent at the beginning of the experiment. The protocol was approved by the Ethics Committee of the Université catholique de Louvain (UCLouvain), Brussels, Belgium. Data presented here were also used (for a different purpose) in another paper (Derosiere et al. 2021b).

### Tokens task

Subjects were seated in front of a computer screen, positioned at a distance of 70 cm from their eyes. Both forearms were placed on the surface of a table with the left and right index fingers placed on a keyboard turned upside down (Fig. 1.A). Subjects performed a variant of the “tokens task” (Cisek et al. 2009, Thura and Cisek 2014, Derosiere et al. 2021a) which was implemented by means of LabView 8.2 (National Instruments, Austin,TX). In this decision-making task, participants had to continuously monitor the distribution of 15 tokens jumping one by one from a central circle to one of two lateral circles. The subjects were instructed to guess which lateral circle would ultimately receive the majority of the tokens; they had to indicate their choice before the last token jump, by pressing a key with the left or right index finger (i.e. a F12 or F5 key-press for the left or right circle, respectively).

**Figure 1:**
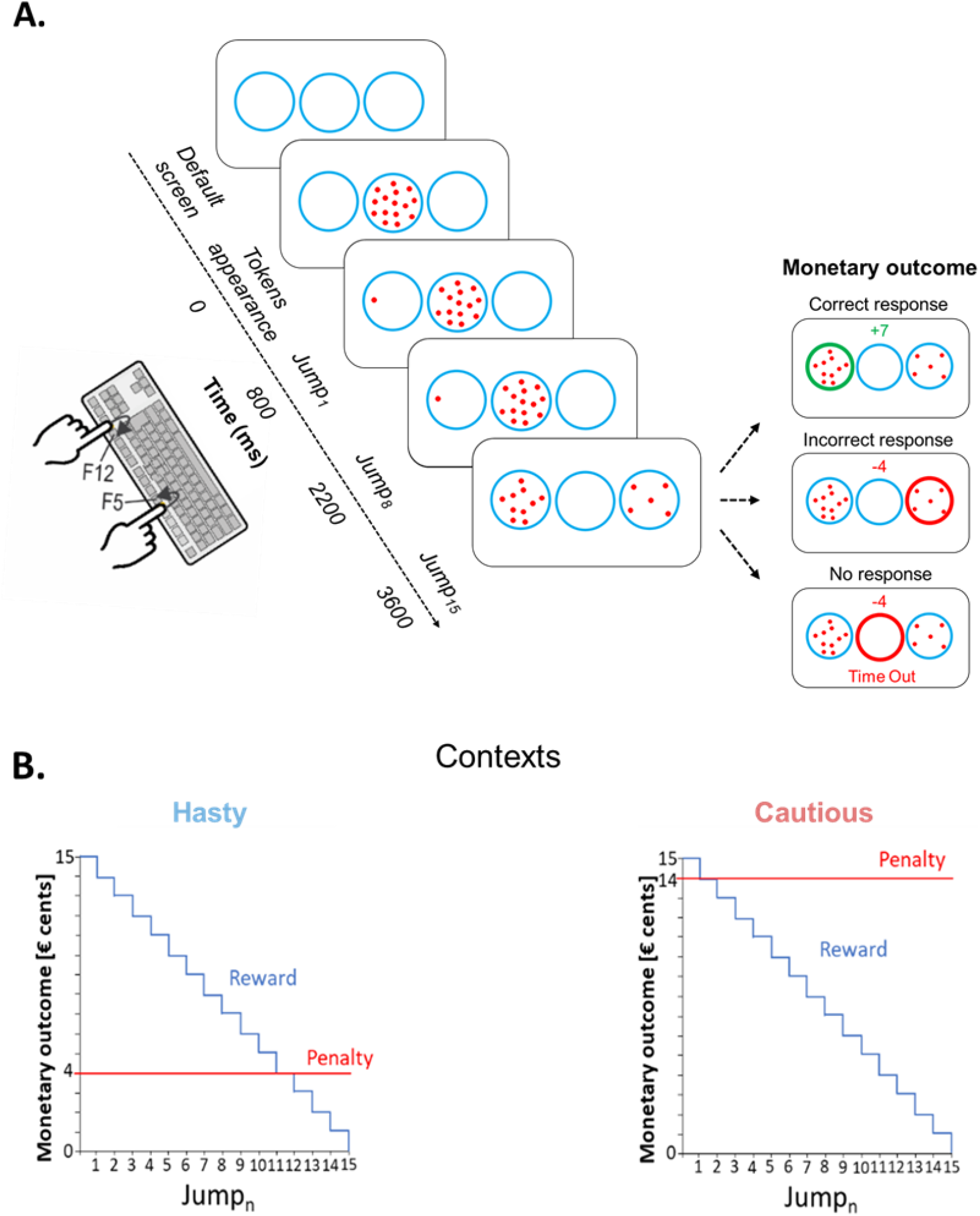
A. Schematic of the tokens task. In each trial, 15 tokens jumped one-by-one every 200 ms from the central circle to one of the lateral circles. The subjects had to indicate by a left or right index finger keypress (i.e., F12 and F5 keys, respectively) which lateral circle they thought would receive the majority of tokens at the end of the trial. For a correct response, the subjects won, in € cents, the number of tokens remaining in the central circle at the time of the response. Hence, the reward earned for a correct response decreased over time, as depicted in B. The right side of panel A depicts the monetary outcome in three exemplary cases. The upper inset represents the reward provided for a correct response between Jump_8_ and Jump_9_, that is when 7 tokens remain in the central circle at the moment the left circle is chosen; the middle inset represents the penalty for an incorrect response in the hasty context, fixed at -4 cents; the lower inset shows the penalty in a “Time Out” trial (no response), fixed at -4 cents, regardless of the context. For representative purposes, the “Time Out” message is depicted below the circles in this example, while it was presented on top of the screen in the actual experiment. **B. Contexts**. Incorrect responses led to a fixed negative score, which differed depending on the context. In the hasty context (shown on the left), the penalty was low, equaling only 4 cents (see red line), promoting fast decisions. In contrast, in the cautious context (shown on the right), the penalty was high, equaling 14 cents, promoting thus slower decisions.

As depicted on Figure 1.A., in between trials, subjects were always presented with a default screen, consisting of three blue circles (4.5 cm diameter each) displayed on a white background for 2500 ms. Each trial started with the appearance of the 15 tokens randomly arranged in the central circle. After a delay of 800 ms, a first token jumped towards the left or right circle, followed every 200 ms, by the other tokens, jumping one by one, to one of the two lateral circles. Subjects were asked to respond as soon as they felt sufficiently confident. The reaction time (RT) was calculated by computing the difference between the time at which subjects pressed the key to indicate their choice and the time of the first tokens jump (Jump_1_). After subjects had pressed the corresponding key, the tokens kept jumping every 200 ms until the central circle was empty (i.e. 2800 ms after Jump_1_). So, the feedback appeared only once all tokens were distributed. At this time, the chosen circle was highlighted either in green or in red depending on whether the response was correct or not, respectively. In addition, a numerical score displayed above the central circle provided subjects with a feedback of their performance (see the ‘Reward, penalty and SAT contexts’ section below). In the absence of any response before the last jump, the central circle turned red with a “Time Out” message and a “-4” (score) appeared on top of the screen. The feedback screen lasted for 500 ms and then disappeared at the same time as the tokens did (the circles always remained on the screen), denoting the end of the trial. From the appearance of the tokens in the central circle, each trial lasted for 6600 ms.

One key feature of the tokens task is that it allows one to calculate, in each trial, the “success probability” *p*_*i*_*(t)* associated with choosing the correct circle *i* at each moment in time *t*. For example, for a total of 15 tokens, if at a particular moment in time the right *(R)* circle contains *NR* tokens, the left *(L)* circle contains *NL* tokens, and the central *(C)* circle contains *NC* tokens, then the probability that the circle on the left will ultimately be the correct one (i.e. the success probability of guessing left) is described as follows:

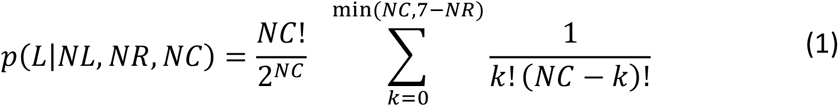

Although the token jumps appeared completely random to subjects, the direction of each jump was determined a priori, producing different types of trials according to specific temporal profiles of pi(t). There were four trial types: ambiguous, obvious, misleading, and arbitrary. The majority of trials (60 %) were ambiguous, as the initial jumps were balanced between the lateral circles, keeping the p_i_(t) close to 0.5 until late in the trial (i.e., p_i_(t) remained between 0.5 and 0.66 up to the Jump_10_). 15 % of trials were “obvious”, meaning that the initial token jumps consistently favored the correct circle (i.e., p_i_(t) was already above 0.7 after Jump_3_ and above 0.8 after jump_5_). 15% of the trials were “misleading”, where most of the first token jumps occurred towards the incorrect lateral circle (i.e., p_i_(t) remained systematically below 0.4 until Jump_3_; from then on, the following tokens jumped mainly in the other direction, that is, towards the circle that eventually turned out being correct). Finally, we included 10 % of trials that were completely arbitrary. These different types of trials were always presented in a randomized order.

### Reward, penalty and SAT contexts

As mentioned above, at the end of each trial, subjects received a feedback score. Correct responses led to a positive score (i.e., a reward) while incorrect responses led to a negative score (i.e., a penalty). Subjects were told that the sum of these scores would turn into a monetary reward at the end of the experiment.

In correct trials, the reward corresponded to the number of tokens remaining in the central circle at the time of the response (in € cents). Hence, reward for a correct choice in a given trial gradually decreased over time (Fig. 1.B.). For instance, a correct response provided between Jump_5_ and Jump_6_ led to a gain of 10 cents (10 tokens remaining in the central circle). However, it only led to a gain of 5 cents when the response was provided between Jump_10_ and Jump_11_ (5 tokens remaining in the central circle). Hence, using a reward dropping over time increased time pressure over the course of a trial and pushed subjects to respond as fast as possible (Derosiere et al. 2019, Derosiere et al. 2021b).

The penalty provided for incorrect choices did not depend on the time taken to choose a lateral circle. Importantly though, it differed between two contexts., In a first context, the cost of making an incorrect choice was low as the penalty was only of -4 cents, pushing subjects to make hasty decisions in order to get high reward scores (hasty context). Conversely, incorrect choices were severely sanctioned in the second context as the penalty there was of -14 cents, emphasizing the need for cautiousness (cautious context).

Moreover, not providing a response before Jump_15_ (i.e., time out trials) also led to a penalty, which was of -4 cents both in the hasty and in the cautious contexts. Hence, in the hasty context, providing an incorrect response or not responding led to the same penalty (i.e., -4 cents), further increasing the urge to respond before the end of the trial in this context. Conversely, in the cautious context, the penalty for making an incorrect choice was much higher than that obtained for an absence of response (i.e., -14 vs -4 cents, respectively), further increasing subjects’ cautiousness in this context.

Hence, with these two contexts, we could consider post-error behavioral adjustments depending on whether the cost of errors was either low or high, prompting the subjects to put the emphasis on decision speed (low accuracy) or on decision accuracy (high accuracy), respectively. As mentioned above, we expected to observe a post-error slowing (PES) in both cases but predicted that it would be more adaptive in the hasty than in the cautious blocks.

### Sensory evidence at RT

The tokens task also allowed us to assess the amount of sensory evidence (i.e. available information) supporting the subjects’ choice at the RT. To estimate the level of sensory evidence at RT, we computed a first-order estimation as the sum of log-likelihood ratios (SumLogLR) of individual token movements at this time (Cisek et al.,2009):

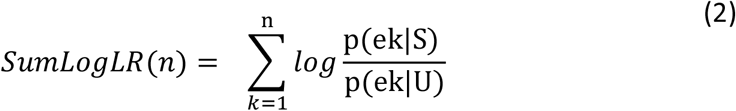

In this equation, p(e_k_|S) is the likelihood of a token event e_k_ (a token jumping into either the chosen or unchosen lateral circle) during trials in which the chosen circle S is correct, and p(e_k_|NS) is its likelihood during trials in which the unchosen circle NS is correct. The SumLogLR is proportional to the difference between the number of tokens contained in each lateral circle; the larger the number of tokens in the chosen circle, as compared to the unchosen circle, the higher is the evidence for the choice and thus the SumLogLR (Derosiere et al. 2019). We expected the latter to be overall higher in the cautious than in the hasty context, reflecting the higher evidence needed before committing to an accurate choice in the former context (Ratcliff 2002, Heitz 2014, Miletić et al. 2021).

### Experimental procedure

Subjects performed the task in the two contexts in two different experimental sessions conducted on separate days at a 24-h interval. The order of the two sessions (i.e. hasty and cautious) was counterbalanced across participants. As described below, each session involved the same structure, except for the addition of a familiarization block in the first session only, to allow subjects to become acquainted with the basic principles of the task (this was of course not necessary for the session coming on the second day).

Each session started with two short blocks involving a simple reaction time (SRT) task. This task was similar to the tokens task described above except that, here, all tokens jumped simultaneously into one of the two lateral circles. The subjects were instructed to respond as fast as possible by pressing the appropriate key (i.e. F12 or F5 for the left or the right circle, respectively). In a given SRT block, the tokens jumped always into the same circle, and subjects were informed in advance of the circle to choose within a block. This SRT task allowed us to estimate the sum of the delays attributable to the sensory and motor processes in the absence of a choice, as achieved in past studies (Cisek et al. 2009, Thura et al. 2014).

Then, subjects performed a few practice blocks. The first one (10 trials) consisted in a version of the tokens task in which the feedback was simplified; indicating only if the subjects’ choice was correct or incorrect by highlighting the chosen circle in green or red, respectively; no reward or penalty was provided here. This first practice block served to familiarize subjects with the basic aspects of the task and was only used during the first session. The practice then continued with two blocks (20 trials each) where subjects performed the task in the context they would be involved in for the whole session (hasty or cautious blocks).

After that, the actual experiment involved 8 blocks of 40 trials (320 trials per session; 640 trials per subject). Each block lasted about 4 min and a break of 2 to 5 minutes was provided between each block. Each session lasted approximatively 150 minutes.

### Statistical analyses

The analyses comprised two parts: first, we ran some tests to check that our manipulation of the penalty indeed led the subject to adopt different SAT policies in the two contexts. Second, and more related to the goal of the current study, we performed analyses to compare the post-error adjustments in the two contexts. Most of the statistical comparisons involved repeated-measures analyses of variance (ANOVA_RM_) run with the Statistica software (version 10.0, Statsoft, Oklahoma United-States). Post-hoc comparisons were conducted using the Tukey’s Honestly Significant Difference (HSD) procedure. The significance level was set at p < .05. Moreover, for the analyses regarding post-error adjustments, we ran a Bayesian equivalent of the ANOVA_RM_ (and t-tests) with JASP (Wagenmakers et al. 2018). In this case, the Bayes Factor (BF_10_) quantifies the evidence for the alternative hypothesis against the null hypothesis. All data are presented as mean ± SE.

#### Manipulation check

In order to verify that our manipulation of penalty (−4 or -14 cents) successfully induced SAT adaptations, we considered the RT, the percentage of correct choices (%Correct) and the SumLogLR at RT in the two contexts. Overall, we expected to observe larger values for these variables in the high penalty (−14 cents) than in the low penalty (−4 cents) blocks, supporting a more conservative behavior in the cautious context compared to the hasty one. To address this directly, we analyzed each variable using two-way ANOVAs_RM_ with CONTEXT (hasty or cautious) and TRIAL_TYPE (obvious, ambiguous or misleading) as within-subject factors.

#### Post-error adjustments

All analyses on post-error adjustments focused on behavior in ambiguous trials. This allowed us to characterize post-error adjustments in a homogeneous set of (ambiguous) trials. We investigated behavior in these trials, referred to as the “n” trials (trials_n_), according to whether they followed an error or a correct choice. These trials preceding trials_n_ are referred to as trial_n-1_ and were separated according to whether they were ambiguous or misleading; there were too few errors in obvious trials to consider them as trials_n-1_. Thus, we considered post-error adjustments on ambiguous trials according to the nature of trials_n-1_ (ambiguous or misleading). For this analysis, we had to exclude 14 participants who had less than five trials_n_ in at least one of the experimental conditions. As a result, statistical analyses were run on a total of 29 subjects (17 women: 23.4 ± 2.4 years old).

There are different methods for quantifying post-error adjustments in trials_n_ (Hajcak and Simons 2002, Dutilh et al. 2012). In the present study, we used a traditional approach consisting in calculating deltas (Δ) for the RT (ΔRT) and for the %Correct (Δ%Correct) in trials_n_ as follows: ΔRT was obtained by calculating the difference between the RT in correct trials_n_ that either followed an error or a correct choice in trials_n-1_ (Williams et al. 2016, Damaso et al. 2020, Smith et al. 2020). Similarly, Δ%Correct corresponded to the difference in %Correct between trials_n_ following an error or a correct choice in trials_n-1_. Hence, a PES manifests as a positive ΔRT. If this positive ΔRT is associated with a positive Δ%Correct, it means that the PES is adaptive (i.e. is associated with a gain in decision accuracy) while a null or negative Δ%Correct reflects a maladaptive PES (no gain or drop in decision accuracy). These ΔRT and Δ%Correct were analyzed using two-ways ANOVA_RM_ with CONTEXT (hasty or cautious) and trials_n-1_-TYPE (ambiguous or misleading) as within-subject factors.

## 3) Results

### Manipulation check

On average, subjects displayed RTs of 1866 ± 457 ms; they performed with a %Correct of 81 ± 18 %, and did so for a level of evidence corresponding to 0.35 ± 0.72 (SumLogLR at RT value; a.u.). Importantly, as depicted in Figure 2 (upper panel), all these values were lower when the penalty was low (i.e. equal to -4 cents) compared to when it was high (i.e. equal to -14 cents), supporting a shift from a cautious to a hasty response policy when the penalty was low (all CONTEXT F_1,28_ > 7.3, all p < .05, see Table 1).

**Table 1.**
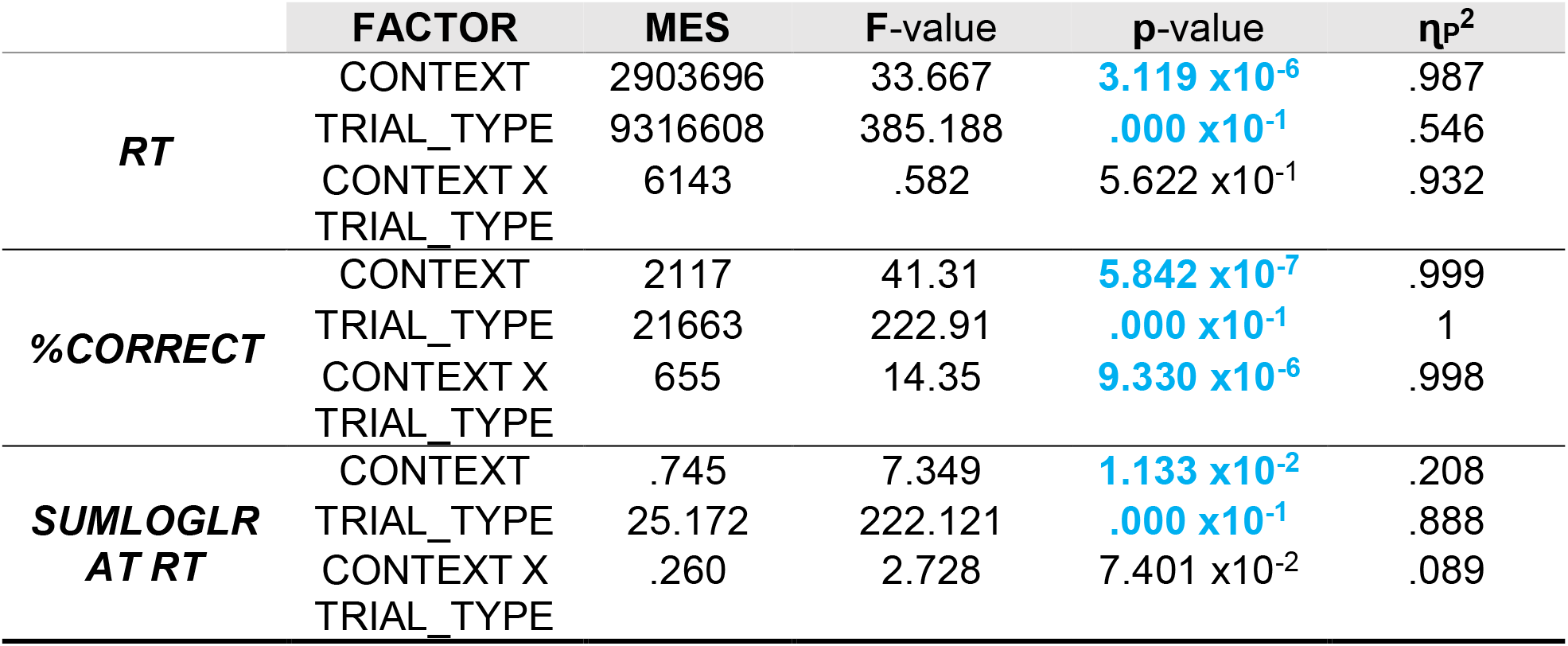
Inferential analyses of behavioral adaptations to the SAT context. The main error square (MES), critical F-value, p-value and partial eta-squared (η_P_^2^) are provided for each factor (CONTEXT; TRIAL_TYPE) and their interaction (CONTEXT X TRIAL_TYPE), following the analysis of reaction time (RT), percentage of correct choices (%Correct) and sensory evidence at RT (SumLogLR at RT). Significant p-values are highlighted in bold and blue.

| | FACTOR | MES | F-value | p-value | $\eta_p^2$ |
| --- | --- | --- | --- | --- | --- |
| **RT** | CONTEXT | 2903696 | 33.667 | **$3.119 \times 10^{-6}$** | .987 |
|  | TRIAL_TYPE | 9316608 | 385.188 | **$.000 \times 10^{-1}$** | .546 |
| | CONTEXT X TRIAL_TYPE | 6143 | .582 | $5.622 \times 10^{-1}$ | .932 |
| **%CORRECT** | CONTEXT | 2117 | 41.31 | **$5.842 \times 10^{-7}$** | .999 |
|  | TRIAL_TYPE | 21663 | 222.91 | **$.000 \times 10^{-1}$** | 1 |
|  | CONTEXT X TRIAL_TYPE | 655 | 14.35 | **$9.330 \times 10^{-6}$** | .998 |
| **SUMLOGLR AT RT** | CONTEXT | .745 | 7.349 | **$1.133 \times 10^{-2}$** | .208 |
|  | TRIAL_TYPE | 25.172 | 222.121 | **$.000 \times 10^{-1}$** | .888 |
| | CONTEXT X TRIAL_TYPE | .260 | 2.728 | $7.401 \times 10^{-2}$ | .089 |

**Figure 2:**
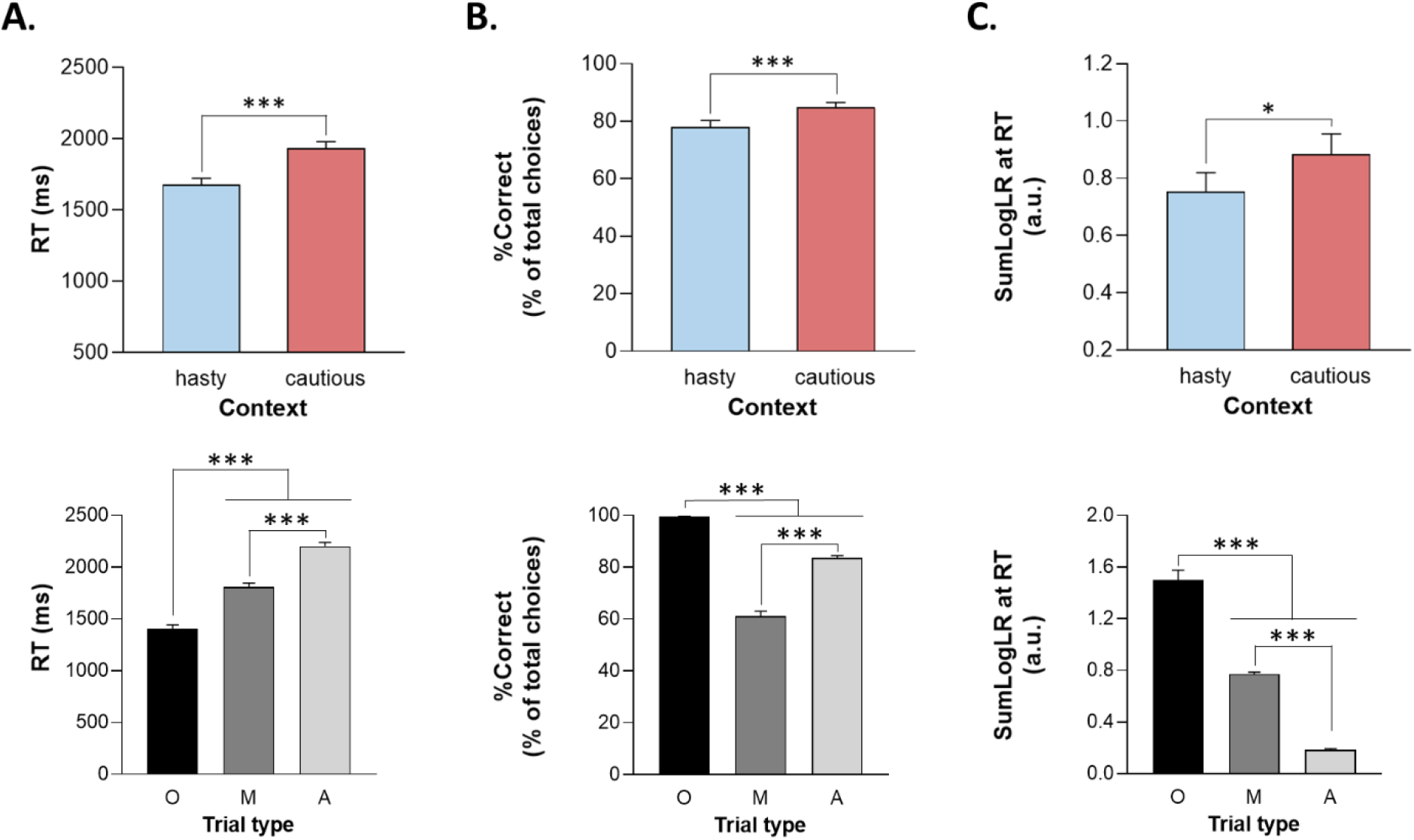
Reaction time (A; RT), percentage of correct choices (B; %Correct) and sensory evidence at RT (C; SumLogLR at RT), depending on the context (upper panel; hasty or cautious) and the trial type (lower panel; Obvious (O), Misleading (M) or Ambiguous (A)). Error bars represent SE. *p < .05, ***p < .001: significantly different.

In addition, as shown on the lower panel of Figure 2, the ANOVA_RM_ revealed an effect of the TRIAL_TYPE on all three parameters (all F_2,56_ > 222, p < .001). As expected, subjects responded faster and more accurately in the obvious trials than in the other trials (all p < .001). They were also faster in misleading than in ambiguous trials (p < .001) but showed a lower accuracy (i.e. lower %Correct) in the former trial type (p < .001), consistent with their misleading nature. Regarding the SumLogLR at RT, it was the highest in the obvious and the lowest in the ambiguous trials (all p < .001), consistent with the different predefined pattern of token jumps in these different trial types.

Finally, the RT and SumLogLR at RT did not display any significant CONTEXT x TRIAL_TYPE interaction (all F_2,56_ < 2.8, all p > .05). Yet, as depicted in Figure 3, this interaction was significant for the %Correct (F_2,56 =_ 14.35, p < .001). As such, the %Correct was larger in the cautious context relative to the hasty one but only in ambiguous and misleading trials (p < .01 and p < .001, respectively). In fact, the obvious trials were so easy that subjects did not make mistakes in this trial type whatever the context.

**Figure 3:**
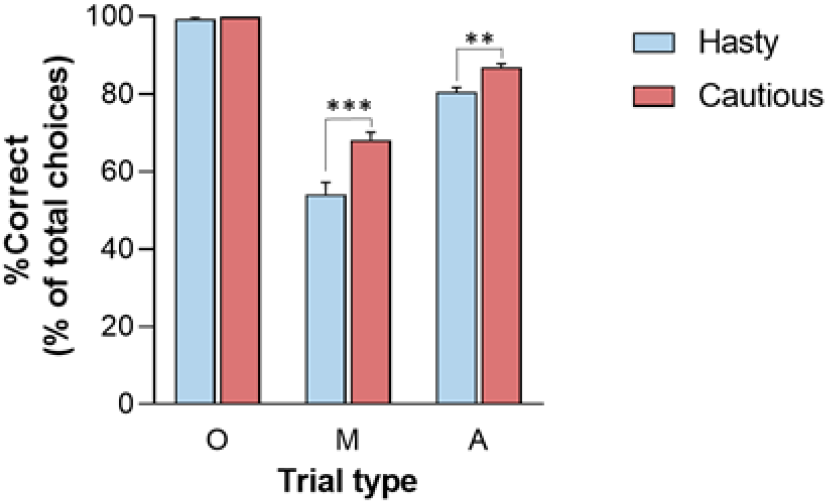
CONTEXT x TRIAL_TYPE interaction on percentage of correct choices (%Correct). %Correct was lower in the hasty (blue bars) than in the cautious (red bars) context when considering Misleading (M) and Ambiguous (A) trials but not for the Obvious (O) trials. Note the absence of errors in these latter trials (%Correct=100), whether in the hasty or cautious context. **p < .01, ***p < .001: significantly different.

### Post-error adjustments

Post-error adjustments (ΔRT and Δ%Correct), calculated with the traditional approach (Dutilh et al. 2012), are displayed in Figure 4 for trials_n_ (always ambiguous), following either ambiguous or misleading trials_n-1_. Even if ΔRT values were positive in all conditions, which would be consistent with the occurrence of a PES, Student’s t-tests against 0 showed that this slowdown was only significant in the hasty context (ΔRT significantly above 0 with a Bonferroni-corrected threshold of .05/4), regardless of the TRIAL_n-1__TYPE (both t_29_ > 3, p = [.0003 .005], Cohen’s *d* = [.567 .767]); t-tests did not reveal any significant difference between ΔRT and 0 in the cautious context (both TRIAL_n-1__TYPE t_29_ > .8, p = [.024 .401], Cohen’s *d* = [.158 .444]). Similarly, equivalent Bayesian analyses (Wagenmakers et al. 2018) showed moderate to strong evidence for a PES in the speed context (BF_10_ = [8.330 98.677]), and moderate evidence for a lack of adjustment after an error in the caution context (BF_10_ = [.275 2.207]). Consistently, the ANOVA_RM_ revealed a significant effect of CONTEXT on ΔRT (F_1,28_ = 6.26, p = .018), in the absence of TRIAL_n-1__TYPE effect (F_1,28_ = .49, p = .49) or CONTEXT x TRIAL_n-1__TYPE interaction (F_1,28_ = 1.39, p = .25). These results were supported by a Bayesian analysis showing moderate evidence for a context effect (BF_10_ = 5.411, see Table 2).

**Table 2.** Inferential analyses of behavioral changes in trial_n_. The prior inclusion probability P(incl), the posterior inclusion probability P(incl|data) and the change from prior to posterior inclusion odds (BF_incl_) are provided for each factor (CONTEXT; TRIAL_n-1__TYPE) and their interaction (CONTEXT X TRIAL_n-1__TYPE), following the analysis of reaction time and %Correct change in trial_n_ (ΔRT and Δ%Correct). In addition, the BF_10_ grades the strength of evidence for the alternative hypothesis against the null hypothesis and the partial eta-squared (η_P_^2^) represents a measure of the effect size. BF_10_ revealing a significant factor effect are highlighted in bold and blue.

| | FACTOR | P(incl) | P(incl\|data) | BF<sub>incl</sub> | BF<sub>10</sub> | $\eta_p^2$ |
| --- | --- | --- | --- | --- | --- | --- |
| $\Delta RT$ | CONTEXT | .600 | .853 | 3.854 | **5.411** | .183 |
|  | TRIAL<sub>n-1</sub>_TYPE | .600 | .247 | .219 | .255 | .17 |
|  | CONTEXT X TRIAL<sub>n-1</sub>_TYPE | .200 | .065 | .278 | .553 | .05 |
| $\Delta \%Correct$ | CONTEXT | .600 | .322 | .317 | .348 | .051 |
|  | TRIAL<sub>n-1</sub>_TYPE | .600 | .626 | 1.116 | **1.444** | .112 |
|  | CONTEXT X TRIAL<sub>n-1</sub>_TYPE | .200 | .078 | .341 | .283 | .050 |

**Figure 4:**
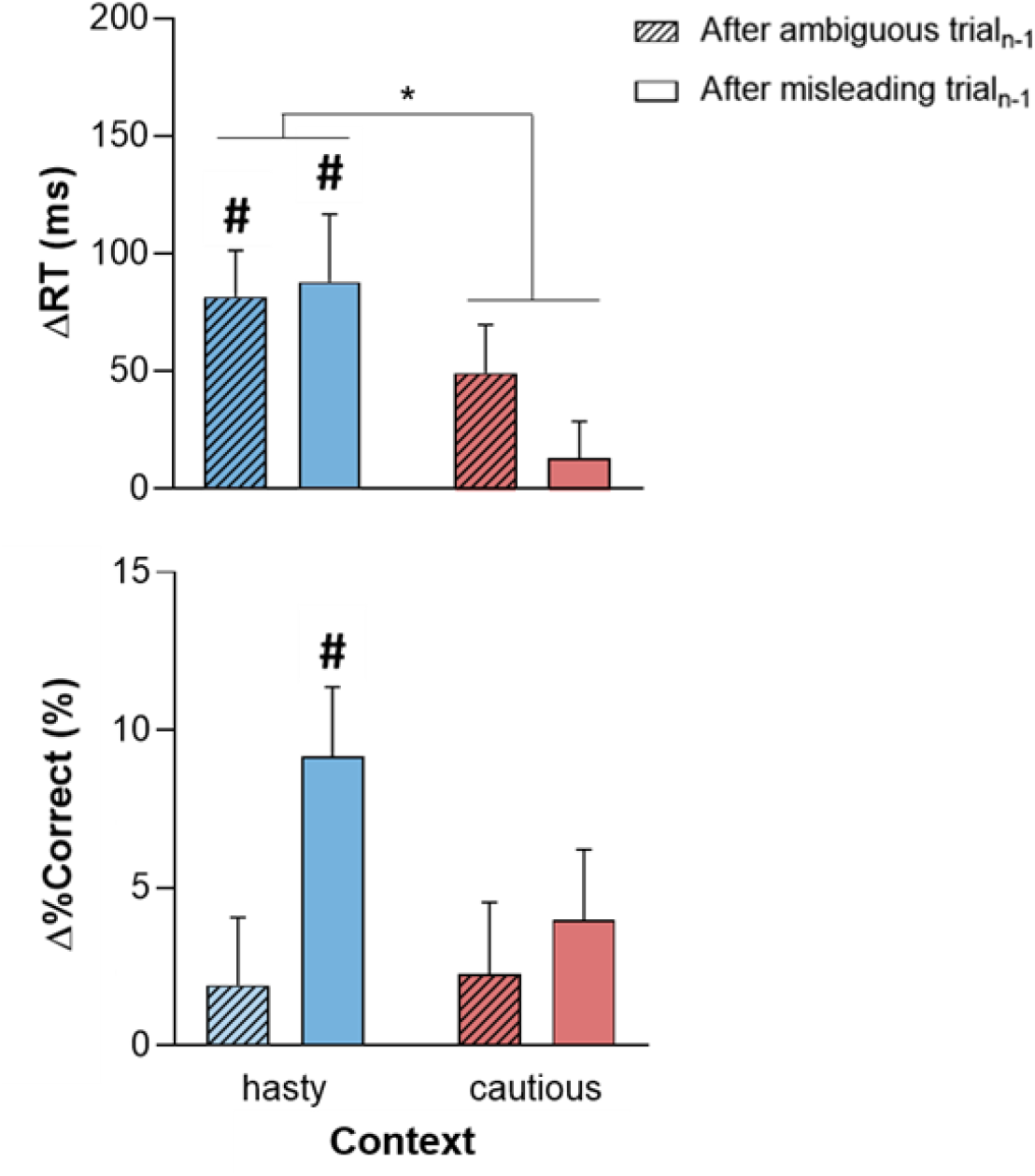
Post-error adjustments of reaction time (ΔRT; upper panel) and %Correct (Δ%Correct; lower panel) depending on the context (hasty or cautious) and on whether trial_n-1_ was ambiguous (crosshatched bars) or misleading (empty bars). While the positive ΔRT in all conditions suggest the presence of a PES, this slowing down was only significant in the hasty context. The latter PES came with a positive Δ%Correct but this effect was only significant following misleading trials_n-1_. Error bars represent SE. #: t-test against 0 (significant difference from 0). *p < .05: significantly different.

Figure 4 (lower panel) evokes positive Δ%Correct in all conditions, which would indicate an increase in decision accuracy in trial_n_. Yet, the Student’s t-tests showed that, this effect was only significant in the hasty context; more surprisingly, it was only present following misleading trials_n-1_ (t_29_ = 4.21, p < .001, Cohen’s *d* = .781 and BF_10_ = 118.923; Bonferroni-corrected threshold= .05/4, see Table 3 for more details). Note though that the variations in Δ%Correct between the different conditions were rather weak, as confirmed by the ANOVA_RM_ analyses which only revealed a marginal effect of TRIAL_n-1__TYPE (F_1,28_ = 3.53, p = .07), with no effect of CONTEXT (F_1,28_ = 1.51, p = .23) or CONTEXT x TRIAL_n-1__TYPE interaction (F_1,28_ = 1.48, p = .23 and BF_10_ = .283). These results were supported by a Bayesian analysis showing anecdotal evidence for a TRIAL_n-1__TYPE effect (BF_10_ = 1.444, see Table 2).

**Table 3.** Inferential Student’s t-tests of behavioral changes in trial_n_. The critical t-value, the p-value and the Cohen’s *d* as a measure of the effect size are represented for each factor (after ambiguous or misleading trials_n-1_ in hasty or cautious context), following the analysis of reaction time and %Correct change in trial_n_ (ΔRT and Δ%Correct). Significant p-value (with a Bonferroni-corrected threshold of .05/4) are highlighted in bold and blue.

|  | FACTOR | t-value | p-value | Cohen's d | BF<sub>10</sub> |
| --- | --- | --- | --- | --- | --- |
| $\Delta RT$ | after ambiguous trials<sub>n-1</sub> in hasty context | 4.131 | **$2.957 \times 10^{-4}$** | .767 | 98.677 |
|  | after misleading trials<sub>n-1</sub> in hasty context | 3.053 | **.005** | .567 | 8.330 |
|  | after ambiguous trials<sub>n-1</sub> in cautious context | 2.390 | .024 | .444 | 2.207 |
|  | after misleading trials<sub>n-1</sub> in cautious context | .853 | .401 | .158 | .275 |
| $\Delta \%Correct$ | after ambiguous trials<sub>n-1</sub> in hasty context | .894 | .379 | .166 | .284 |
|  | after misleading trials<sub>n-1</sub> in hasty context | 4.208 | **$2.4 \times 10^{-4}$** | .781 | 118.923 |
|  | after ambiguous trials<sub>n-1</sub> in cautious context | .994 | .329 | .185 | .310 |
|  | after misleading trials<sub>n-1</sub> in cautious context | 1.772 | .087 | .329 | .786 |

In conclusion, our data indicate that post-error adjustments varied according to the context in which subjects performed the tokens task, with PES being only significant in the hasty context, and a gain in performance being only observed after errors in misleading trials.

## 4) Discussion

Here, we investigated if the nature of post-error adjustments can vary according to whether a subject favors hasty or cautious decisions. To address this point, we had subjects perform the tokens task in separate blocks where errors were either poorly penalized, encouraging hasty responses (but low accuracy), or highly penalized, calling for more cautiousness (at the cost of speed). The results show that, overall, subjects slowed down after erroneous choices, supporting the presence of a PES. Yet, despite the fact that ΔRT values were numerically positive in all conditions, this PES was only significant in the hasty context (after correction for multiple comparisons). Moreover, consistent with an adaptive adjustment in this context, we observed a significant improvement in performance, but only following misleading trials_n-1_; the positive Δ%Correct did not reach significance following ambiguous trials_n-1_, suggesting a maladaptive slow down with no performance gain following this type of trial in the hasty context.

The positive values of ΔRT in all conditions indicate that, if anything, subjects slowed down after an error. However, contrary to our expectation to observe PES in the two contexts, this ΔRT was only significant in the hasty context suggesting that subjects only slowed down when they were in a context emphasizing speed (low accuracy) but not when the context promoted more accurate choices. PES, as observed in the hasty context, is usually associated with a cognitive control process recruited to prevent future errors (Smith and Brewer 1995, Siegert et al. 2014, Beatty et al. 2020). Such process is thought to operate at least in part at the level of the decision threshold, increasing its height with respect to baseline activity as a means to augment the amount of (neural) evidence accumulation required to reach the decision threshold (Dutilh et al. 2012, Purcell and Kiani 2016, Schiffler et al. 2017, Fischer et al. 2018, Alamia et al. 2019, Derosiere et al. 2018, Derosiere et al. 2019, Derosiere et al. 2021b); this of course prolongs the decision time but increases the probability of choosing the right circle and therefore the reward rate.

Consistent with the occurrence of such adaptive adjustment maximizing the reward rate in the hasty context, the PES observed there was associated with positive Δ%Correct values (Botvinick and Braver 2015, Thura 2020, Vassiliadis and Derosiere 2020). Yet surprisingly, this was only true after misleading trial_n-1_ but not after ambiguous trial_n-1_, as Δ%Correct did not reach significance following the latter trial type. Hence, PES in the hasty context was not always associated with a gain in decision accuracy. Rather, it seemed to switch from being adaptive (it helped increase accuracy following misleading trial_n-1_) to maladaptive as it led to no gain in performance after ambiguous trial_n-1_. Such a finding suggests that errors did not solely trigger shifts in decision thresholds. Indeed, if this was the case, one would expect PES to be accompanied by an increase in accuracy regardless of the type of trial_n-1_ in which an error occurred. Alternatively, a non-exclusive possibility is that task engagement varied following errors in these two trial_n-1_ types. We believe this may be the case because post-error task engagement (or arousal) has been shown to vary with the level of confidence at the moment an error is made (Yeung and Summerfield 2012, Purcell and Kiani 2016, Desender et al. 2019), which itself depends on the amount of sensory evidence available to make the (incorrect) choice (Meyniel et al. 2015, Pouget et al. 2016, Sanders et al. 2016, Urai et al. 2017, Desender et al. 2019). Accordingly, past studies have shown that when errors are made based on poor sensory evidence (i.e. with a low confidence level, as in ambiguous trials), arousal decreases significantly in the following trial (Notebaert et al. 2009, Nunez Castellar et al. 2010, Navarro-Cebrian et al. 2013, Wessel 2018, Desender et al. 2019, Purcell and Kiani 2016). By contrast, when errors are related to the presence of high (conflicting) sensory evidence (as in misleading trials), arousal is found increased in the following trial, an effect that may help dedicate attention to relevant sensory evidence (King et al. 2010, Danielmeier and Ullsperger 2011). Hence, it is plausible that in the current study, post-error task engagement was larger following misleading than ambiguous trial_n-1_, allowing PES to result in a performance gain following the former but not the latter trial type. Such a hypothesis could be tested in future work by investigating changes in pupil diameter following errors in our task (Kahneman and Beatty 1966, Saderi et al. 2021).

Critically, Wessel proposed that PES arise from a sequence of processes including first a transient automatic response to the unexpected event (i.e. error), which triggers a reorientation of attention, followed by an adaptive process increasing decision threshold to prevent future errors (Wessel 2018). Based on this adaptive orienting theory, the delay between the feedback on trial_n-1_ (indicating an error) and the start of trial_n_, which corresponds to the intertrial interval (ITI) duration, can influence the nature of PES and needs to be long enough to allow the second adaptive process to take place. This was the case in our study where the ITI duration was 2500 ms which, based on Wessel, is long enough for the adaptive process to occur. Hence, because the ITI duration was also comparable between the PES conditions, it is unlikely that this aspect of the task affected our data.

Unexpectedly, in this study we observed a PES only in the hasty but not in the cautious context. One tempting explanation is that slowing down after errors could only effectively increase accuracy and thus be adaptive in the hasty but not in the cautious context. That is, because subjects emphasized speed in the hasty context, it is likely that a great proportion of errors were made because subjects responded too fast and not necessarily because the trial was difficult (Damaso et al. 2020). Hence, in this context, errors could easily be avoided by slowing down a bit in the following trial. In contrast, subjects were generally more cautious in the other context and it is thus plausible that errors occurred when choices were complex rather than because responses were too hasty (Ratcliff and Rouder 1998, Brown and Heathcote 2008, Ratcliff and McKoon 2008). Slowing down following these trials may not be effective as it would not necessarily enhance accuracy; that is, even if subjects fail on the most complex choices, they are generally cautious enough to succeed on most trials and slowing down further would not lead to any performance gain. Yet, we believe such an explanation does not hold here. As such, it is important to note that RTs in cautious blocks were around 2017 ms, which falls between Jump_10_ and Jump_11_, coinciding thus with the moment sensory evidence in favor of the correct choice starts to increase greatly (see methods). This means that even if subjects were already generally cautious (and slower) in this context, slowing down would have been adaptive because it would have allowed to provide responses based on more evidence.

A more plausible explanation is that the absence of PES in the cautious context is related to the way we promoted cautiousness in the current study. As such, changes in the SAT policy between the two contexts were engendered by manipulating the penalty size. However, even if error punishment is known to increase cautiousness (Potts 2011, Derosiere et al. 2021b), as desired here, monetary losses also generate an emotional response (Carver 2006, Simoes-Franklin et al. 2010, Frijda et al. 2014, Eben et al. 2020a), a sense of frustration increasing with the size of the loss (Gehring et al. 2002, Holroyd et al. 2004, Yeung and Sanfey 2004, Eben et al. 2020c). Importantly, such negative emotion has been shown to induce a post-error acceleration of RTs rather than a slowdown (Purcell and Kiani 2016, Verbruggen et al. 2017, Dyson et al. 2018, Damaso et al. 2020, Eben et al. 2020c, Dyson 2021). Accordingly, several studies have found that subjects act more impulsively after a loss or a nonrewarded trial than a rewarded one (Gipson et al. 2012, Verbruggen et al. 2017, Eben et al. 2020c). Altogether, this literature suggests that the emotional response to monetary loss might have precluded us from observing a PES in the cautious context. In other words, errors in the cautious context may have triggered opposite reactions counteracting each other; that is, a frustration feeling due to the high penalty (speeding up behavior) and an adaptive adjustment to prevent hasty errors (slowing down behavior). In the future, it would be interesting to dissociate the manipulation of the context from that of the penalty. Moreover, as the level of punishment sensitivity impacts error monitoring (Unger et al. 2012, Laurent et al. 2018), it also seems relevant to add some questionnaires to measure this personality trait such as the behavioral inhibition system (BIS) scale.

Interpretation of the current data is limited by the fact that a large number of subjects were excluded from the analyses because of an insufficient number of trials, reducing thus the sample size and the statistical power. In addition, the low error rate in the cautious context and the presence of different trial types impacted also the calculation of PES by preventing the use of another method than the traditional one. We recognize that this traditional method is prone to different biases such as global fluctuations in subject performance or in the number of post-correct trials outnumbering the number of post-error trials (Schroder et al. 2020). Note that even if some studies show that these biases can lead to an underestimation of post-error adjustments by decreasing effect sizes (Damaso et al. 2020, Schroder et al. 2020), others suggest that these biases do not radically change the results (van den Brink et al. 2014, Murphy et al. 2016).

In conclusion, our findings highlight a complex combination of processes that come into play following errors and that affect the speed of ensuing actions as well as the degree to which such post-error adjustment comes with a gain in performance or is rather maladaptive. The recruitment of these processes depends on several factors, including the context within which choices are made and the nature of erroneous trials, which affect altogether the subjects’ strategy, their engagement in the task and likely also their emotional reaction to the error.

## Acknowledgements

This work was supported by the grant from the Belgian National Funds for Scientific Research (FRIA-B2; INHIBACTION). We thank Sara Lo Presti, Roxanne Weverbergh and Caroline Hermand for their help in the acquisition of the data.

## Declaration of interests

The authors declare no competing interests.

## Notes

### Competing Interest Statement

The authors have declared no competing interest.

## REFERENCES

Alamia A, Duque J, VanRullen R, Zenon A, Derosiere G. (2019). “Implicit visual cues tune oscillatory motor activity during decision-making.” NeuroImage 186; 424–436. doi: 10.1016/j.neuroimage.2018.11.027

Beatty, P. J., G. A. Buzzell, D. M. Roberts and C. G. McDonald (2020). “Contrasting time and frequency domains: ERN and induced theta oscillations differentially predict post-error behavior.” Cogn Affect Behav Neurosci 20(3): 636–647. doi:10.3758/s13415-020-00792-7

Botvinick, M. and T. Braver (2015). “Motivation and cognitive control: from behavior to neural mechanism.” Annu Rev Psychol 66: 83–113. doi:10.1146/annurev-psych-010814-015044

Brown, S. D. and A. Heathcote (2008). “The simplest complete model of choice response time: linear ballistic accumulation.” Cogn Psychol 57(3): 153–178. doi:10.1016/j.cogpsych.2007.12.002

Carver, C. S. (2006). “Approach, Avoidance, and the Self-Regulation of Affect and Action.” Motivation and Emotion 30(2): 105–110. doi:10.1007/s11031-006-9044-7

Cavanagh, J. F., J. L. Sanguinetti, J. J. Allen, S. J. Sherman and M. J. Frank (2014). “The subthalamic nucleus contributes to post-error slowing.” J Cogn Neurosci 26(11): 2637–2644. doi:10.1162/jocn_a_00659

Ceccarini, F., S. Guerra, S. Betti, A. Vergazzini, L. Sartori and U. Castiello (2019). “Changes in corticospinal excitability associated with post-error slowing.” Cortex 120: 92–100. doi:https://doi.org/10.1016/j.cortex.2019.05.015

Chittka, L., P. Skorupski and N. E. Raine (2009). “Speed-accuracy tradeoffs in animal decision making.” Trends Ecol Evol 24(7): 400–407. doi:10.1016/j.tree.2009.02.010

Cisek, P., G. A. Puskas and S. El-Murr (2009). “Decisions in changing conditions: the urgency-gating model.” J Neurosci 29(37): 11560–11571. doi:10.1523/JNEUROSCI.1844-09.2009

Compton, R. J., D. Gearinger, H. Wild, D. Rette, E. C. Heaton, S. Histon, P. Thiel and M. Jaskir (2021). “Simultaneous EEG and pupillary evidence for post-error arousal during a speeded performance task.” Eur J Neurosci 53(2): 543–555. doi:10.1111/ejn.14947

Damaso, K., P. Williams and A. Heathcote (2020). “Evidence for different types of errors being associated with different types of post-error changes.” Psychon Bull Rev. doi:10.3758/s13423-019-01675-w

Danielmeier, C. and M. Ullsperger (2011). “Post-error adjustments.” Front Psychol 2: 233. doi:10.3389/fpsyg.2011.00233

Derosiere, G., D. Thura, P. Cisek and J. Duque (2019). “Motor cortex disruption delays motor processes but not deliberation about action choices.” Journal of Neurophysiology 122(4): 1566–1577. doi:10.1152/jn.00163.2019

Derosiere, G., D. Thura, P. Cisek and J. Duque (2021a). “Trading accuracy for speed over the course of a decision.” Journal of Neurophysiology 126(2);361-372. doi: https://doi.org/10.1152/jn.00038.2021

Derosiere, G., D. Thura, P. Cisek and J. Duque (2021b). “Overlapping influences shape motor activity during hasty sensorimotor decisions.” bioRxiv: 2021.2008.2006.455419.

Derosiere G, Klein PA, Nozaradan S, Zenon A, Mouraux A, Duque J. (2018). Visuomotor correlates of conflict expectation in the context of motor decisions. The Journal of Neuroscience 38(44); 9486–9504. doi: 10.1523/JNEUROSCI.0623-18.2018

Desender, K., A. Boldt, T. Verguts and T. H. Donner (2019). “Confidence predicts speed-accuracy tradeoff for subsequent decisions.” Elife 8. doi:10.7554/eLife.43499

Dubravac, M., C. M. Roebers and B. Meier (2020). “Different temporal dynamics after conflicts and errors in children and adults.” PLoS One 15(8): e0238221. doi:10.1371/journal.pone.0238221

Dutilh, G., D. van Ravenzwaaij, S. Nieuwenhuis, H. L. J. van der Maas, B. U. Forstmann and E.-J. Wagenmakers (2012). “How to measure post-error slowing: A confound and a simple solution.” Journal of Mathematical Psychology 56(3): 208–216. doi:https://doi.org/10.1016/j.jmp.2012.04.001

Dutilh, G., J. Vandekerckhove, B. U. Forstmann, E. Keuleers, M. Brysbaert and E.-J. Wagenmakers (2012). “Testing theories of post-error slowing.” Attention, Perception, & Psychophysics 74(2): 454–465. doi:10.3758/s13414-011-0243-2

Dyson, B. J. (2021). “Variability in competitive decision-making speed and quality against exploiting and exploitative opponents.” Sci Rep 11(1): 2859. doi:10.1038/s41598-021-82269-2

Dyson, B. J., J. Sundvall, L. Forder and S. Douglas (2018). “Failure generates impulsivity only when outcomes cannot be controlled.” J Exp Psychol Hum Percept Perform 44(10): 1483–1487. doi:10.1037/xhp0000557

Eben, C., J. Billieux and F. Verbruggen (2020a). “Clarifying the Role of Negative Emotions in the Origin and Control of Impulsive Actions.” Psychologica Belgica 60: 1–17. doi:10.5334/pb.502

Eben, C., Z. Chen, E. Cracco, M. Brass, J. Billieux and F. Verbruggen (2020b). “Are post-error adjustments influenced by beliefs in free will? A failure to replicate Rigoni, Wilquin, Brass and Burle, 2013.” R Soc Open Sci 7(11): 200664. doi:10.1098/rsos.200664

Eben, C., Z. Chen, L. Vermeylen, J. Billieux and F. Verbruggen (2020c). “A direct and conceptual replication of post-loss speeding when gambling.” R Soc Open Sci 7(5): 200090. doi:10.1098/rsos.200090

Fischer, A. G., R. Nigbur, T. A. Klein, C. Danielmeier and M. Ullsperger (2018). “Cortical beta power reflects decision dynamics and uncovers multiple facets of post-error adaptation.” Nat Commun 9(1): 5038. doi:10.1038/s41467-018-07456-8

Frijda, N. H., K. R. Ridderinkhof and E. Rietveld (2014). “Impulsive action: emotional impulses and their control.” Front Psychol 5: 518. doi:10.3389/fpsyg.2014.00518

Fu, Z., D. J. Wu, I. Ross, J. M. Chung, A. N. Mamelak, R. Adolphs and U. Rutishauser (2019). “Single-Neuron Correlates of Error Monitoring and Post-Error Adjustments in Human Medial Frontal Cortex.” Neuron 101(1): 165–177 e165. doi:10.1016/j.neuron.2018.11.016

Gehring William, J. and R. Willoughby Adrian (2002). “The Medial Frontal Cortex and the Rapid Processing of Monetary Gains and Losses.” Science 295(5563): 2279–2282. doi:10.1126/science.1066893

Gipson, C. D., J. S. Beckmann, Z. W. Adams, J. A. Marusich, T. O. Nesland, J. R. Yates, T. H. Kelly and M. T. Bardo (2012). “A translational behavioral model of mood-based impulsivity: Implications for substance abuse.” Drug Alcohol Depend 122(1-2): 93-99. doi:10.1016/j.drugalcdep.2011.09.014

Guo, X., Z. Luo and X. Yu (2019). “A Speed-Accuracy Tradeoff Hierarchical Model Based on Cognitive Experiment.” Front Psychol 10: 2910. doi:10.3389/fpsyg.2019.02910

Hajcak, G., N. McDonald and R. F. Simons (2003). “To err is autonomic: error-related brain potentials, ANS activity, and post-error compensatory behavior.” Psychophysiology 40(6): 895–903. doi:10.1111/1469-8986.00107

Hajcak, G. and R. F. Simons (2002). “Error-related brain activity in obsessive-compulsive undergraduates.” Psychiatry Res 110(1): 63–72. doi:10.1016/s0165-1781(02)00034-3

Heitz, R. P. (2014). “The speed-accuracy tradeoff: history, physiology, methodology, and behavior.” Front Neurosci 8: 150. doi:10.3389/fnins.2014.00150

Henmon, V. A. C. (1911). “The relation of the time of a judgment to its accuracy.” Psychological Review 18(3): 186–201. doi:http://dx.doi.org/10.1037/h0074579

Holroyd, C. B., J. T. Larsen and J. D. Cohen (2004). “Context dependence of the event-related brain potential associated with reward and punishment.” Psychophysiology 41(2): 245–253. doi:10.1111/j.1469-8986.2004.00152.x

Houtman, F., E. N. Castellar and W. Notebaert (2012). “Orienting to errors with and without immediate feedback.” Journal of Cognitive Psychology 24(3): 278–285. doi:10.1080/20445911.2011.617301

Kahneman, D. and J. Beatty (1966). “Pupil Diameter and Load on Memory.” Science 154(3756): 1583–1585. doi:10.1126/science.154.3756.1583

King, J. A., F. M. Korb, D. Y. von Cramon and M. Ullsperger (2010). “Post-error behavioral adjustments are facilitated by activation and suppression of task-relevant and task-irrelevant information processing.” J Neurosci 30(38): 12759–12769. doi:10.1523/JNEUROSCI.3274-10.2010

Kirschner, H., J. Humann, J. Derrfuss, C. Danielmeier and M. Ullsperger (2020). “Neural and behavioral traces of error awareness.” Cognitive, Affective, & Behavioral Neuroscience. doi:10.3758/s13415-020-00838-w

Laurent, R., N. C. van Wouwe, M. Turchan, C. Tolleson, F. Phibbs, E. Bradley, W. van den Wildenberg and S. A. Wylie (2018). “Motivational Sensitivities Linked to Impulsive Motor Errors in Parkinson’s Disease.” J Int Neuropsychol Soc 24(2): 128–138. doi:10.1017/S1355617717000741

Li, Q., Q. Long, N. Hu, Y. Tang and A. Chen (2020). “N-Back Task Training Helps to Improve Post-error Performance.” Front Psychol 11: 370. doi:10.3389/fpsyg.2020.00370

Meyniel, F., M. Sigman and Z. F. Mainen (2015). “Confidence as Bayesian Probability: From Neural Origins to Behavior.” Neuron 88(1): 78–92. doi:10.1016/j.neuron.2015.09.039

Miletic, S., R. J. Boag, A. C. Trutti, N. Stevenson, B. U. Forstmann and A. Heathcote (2021). “A new model of decision processing in instrumental learning tasks.” eLife 10. doi:10.7554/eLife.63055

Murphy, P. R., M. L. van Moort and S. Nieuwenhuis (2016) “The Pupillary Orienting Response Predicts Adaptive Behavioral Adjustment after Errors.” PloS one 11, e0151763. doi: 10.1371/journal.pone.0151763

Navarro-Cebrian, A., R. T. Knight and A. S. Kayser (2013). “Error-monitoring and post-error compensations: dissociation between perceptual failures and motor errors with and without awareness.” J Neurosci 33(30): 12375–12383. doi:10.1523/jneurosci.0447-13.2013

Nigbur, R. and M. Ullsperger (2020). “Funny kittens: Positive mood induced via short video-clips affects error processing but not conflict control.” International Journal of Psychophysiology 147: 147–155.

Notebaert, W., F. Houtman, F. V. Opstal, W. Gevers, W. Fias and T. Verguts (2009). “Post-error slowing: an orienting account.” Cognition 111(2): 275–279. doi:10.1016/j.cognition.2009.02.002

Nunez Castellar, E., S. Kuhn, W. Fias and W. Notebaert (2010). “Outcome expectancy and not accuracy determines posterror slowing: ERP support.” Cogn Affect Behav Neurosci 10(2): 270–278. doi:10.3758/CABN.10.2.270

Oldfield, R. C. (1971). “The assessment and analysis of handedness: The Edinburgh inventory.” Neuropsychologia 9(1): 97–113. doi:https://doi.org/10.1016/0028-3932(71)90067-4

Potts, G. F. (2011). “Impact of reward and punishment motivation on behavior monitoring as indexed by the error-related negativity.” Int J Psychophysiol 81(3): 324–331. doi:10.1016/j.ijpsycho.2011.07.020

Pouget, A., J. Drugowitsch and A. Kepecs (2016). “Confidence and certainty: distinct probabilistic quantities for different goals.” Nat Neurosci 19(3): 366–374. doi:10.1038/nn.4240

Purcell, B. A. and R. Kiani (2016). “Neural Mechanisms of Post-error Adjustments of Decision Policy in Parietal Cortex.” Neuron 89(3): 658–671. doi:10.1016/j.neuron.2015.12.027

Rabbitt, P. M. A. and S. M. Vyas (1970). “An elementary preliminary taxonomy for some errors in laboratory choice RT tasks.” Acta Psychologica 33: 56–76. doi:https://doi.org/10.1016/0001-6918(70)90122-8

Ratcliff, R. (2002). “A diffusion model account of response time and accuracy in a brightness discrimination task: Fitting real data and failing to fit fake but plausible data.” Psychonomic Bulletin & Review 9(2): 278–291. doi:10.3758/BF03196283

Ratcliff, R. and G. McKoon (2008). “The diffusion decision model: theory and data for two-choice decision tasks.” Neural Comput 20(4): 873–922. doi:10.1162/neco.2008.12-06-420

Ratcliff, R. and J. N. Rouder (1998). “Modeling Response Times for Two-Choice Decisions.” Psychological Science 9(5): 347–356. doi:10.1111/1467-9280.00067

Reynaud, A. J., C. Saleri Lunazzi and D. Thura (2020). “Humans sacrifice decision-making for action execution when a demanding control of movement is required.” J Neurophysiol 124(2): 497–509. doi:10.1152/jn.00220.2020

Rinberg, D., A. Koulakov and A. Gelperin (2006). “Speed-accuracy tradeoff in olfaction.” Neuron 51(3): 351–358. doi:10.1016/j.neuron.2006.07.013

Saderi, D., Z. P. Schwartz, C. R. Heller, J. R. Pennington and S. V. David (2021). “Dissociation of task engagement and arousal effects in auditory cortex and midbrain.” eLife 10: e60153. doi:10.7554/eLife.60153

Salinas, E., V. E. Scerra, C. K. Hauser, M. G. Costello and T. R. Stanford (2014). “Decoupling speed and accuracy in an urgent decision-making task reveals multiple contributions to their trade-off.” Front Neurosci 8: 85.

Sanders, J. I., B. Hangya and A. Kepecs (2016). “Signatures of a Statistical Computation in the Human Sense of Confidence.” Neuron 90(3): 499–506. doi:10.1016/j.neuron.2016.03.025

Schiffler, B. C., S. L. Bengtsson and D. Lundqvist (2017). “The Sustained Influence of an Error on Future Decision-Making.” Front Psychol 8: 1077. doi:10.3389/fpsyg.2017.01077

Schroder, H. S., S. Nickels, E. Cardenas, M. Breiger, S. Perlo and D. A. Pizzagalli (2020). “Optimizing assessments of post-error slowing: A neurobehavioral investigation of a flanker task.” Psychophysiology 57(2): e13473. doi:10.1111/psyp.13473

Siegert, S., M. Herrojo Ruiz, C. Brucke, J. Huebl, G. H. Schneider, M. Ullsperger and A. A. Kuhn (2014). “Error signals in the subthalamic nucleus are related to post-error slowing in patients with Parkinson’s disease.” Cortex 60: 103–120. doi:10.1016/j.cortex.2013.12.008

Simoes-Franklin, C., R. Hester, M. Shpaner, J. J. Foxe and H. Garavan (2010). “Executive function and error detection: The effect of motivation on cingulate and ventral striatum activity.” Hum Brain Mapp 31(3): 458–469. doi:10.1002/hbm.20879

Smith, D. M., T. Dykstra, E. Hazeltine and E. H. Schumacher (2020). “Task representation affects the boundaries of behavioral slowing following an error.” Atten Percept Psychophys 82(5): 2315–2326. doi:10.3758/s13414-020-01985-5

Smith, G. A. and N. Brewer (1995). “Slowness and age: speed-accuracy mechanisms.” Psychology and aging 10(2): 238.

Sokolov, E. N. (1963). “Higher Nervous Functions: The Orienting Reflex.” Annual Review of Physiology $V 25(1): 545–580.

Spieser, L., M. Servant, T. Hasbroucq and B. Burle (2017). “Beyond decision! Motor contribution to speed–accuracy trade-off in decision-making.” Psychonomic Bulletin & Review 24(3): 950–956. doi:10.3758/s13423-016-1172-9

Steinhauser, M. and S. K. Andersen (2019). “Rapid adaptive adjustments of selective attention following errors revealed by the time course of steady-state visual evoked potentials.” Neuroimage 186: 83–92. doi:10.1016/j.neuroimage.2018.10.059

Thura, D. (2020). “Decision urgency invigorates movement in humans.” Behav Brain Res 382: 112477. doi:10.1016/j.bbr.2020.112477

Thura, D. and P. Cisek (2014). “Deliberation and commitment in the premotor and primary motor cortex during dynamic decision making.” Neuron 81(6): 1401–1416. doi:10.1016/j.neuron.2014.01.031

Thura, D., I. Cos, J. Trung and P. Cisek (2014). “Context-dependent urgency influences speed-accuracy trade-offs in decision-making and movement execution.” J Neurosci 34(49): 16442–16454. doi:10.1523/jneurosci.0162-14.2014

Topor, M., B. Opitz and H. C. Leonard (2021). “Error-Related Cognitive Control and Behavioral Adaptation Mechanisms in the Context of Motor Functioning and Anxiety.” Front Hum Neurosci 15: 615616. doi:10.3389/fnhum.2021.615616

Unger, K., S. Heintz and J. Kray (2012). “Punishment sensitivity modulates the processing of negative feedback but not error-induced learning.” Front Hum Neurosci 6: 186. doi:10.3389/fnhum.2012.00186

Urai, A. E., A. Braun and T. H. Donner (2017). “Pupil-linked arousal is driven by decision uncertainty and alters serial choice bias.” Nat Commun 8: 14637. doi:10.1038/ncomms14637

van den Brink, R. L., S. C. Wynn and S. Nieuwenhuis (2014). “Post-Error Slowing as a Consequence of Disturbed Low-Frequency Oscillatory Phase Entrainment.” The Journal of Neuroscience 34(33): 11096. doi:10.1523/JNEUROSCI.4991-13.2014

van Driel, J., K. R. Ridderinkhof and M. X. Cohen (2012). “Not All Errors Are Alike: Theta and Alpha EEG Dynamics Relate to Differences in Error-Processing Dynamics.” The Journal of Neuroscience 32(47): 16795–16806. doi:10.1523/jneurosci.0802-12.2012

Vassiliadis, P. and G. Derosiere (2020). “Selecting and Executing Actions for Rewards.” J Neurosci 40(34): 6474–6476. doi:10.1523/jneurosci.1250-20.2020

Verbruggen, F., C. D. Chambers, N. S. Lawrence and I. P. McLaren (2017). “Winning and losing: Effects on impulsive action.” J Exp Psychol Hum Percept Perform 43(1): 147–168. doi:10.1037/xhp0000284

Wagenmakers, E. J., J. Love, M. Marsman, T. Jamil, A. Ly, J. Verhagen, R. Selker, Q. F. Gronau, D. Dropmann, B. Boutin, F. Meerhoff, P. Knight, A. Raj, E. J. van Kesteren, J. van Doorn, M. Smira, S. Epskamp, A. Etz, D. Matzke, T. de Jong, D. van den Bergh, A. Sarafoglou, H. Steingroever, K. Derks, J. N. Rouder and R. D. Morey (2018). “Bayesian inference for psychology. Part II: Example applications with JASP.” Psychon Bull Rev 25(1): 58–76. doi:10.3758/s13423-017-1323-7

Wessel, J. R. (2018). “An adaptive orienting theory of error processing.” Psychophysiology 55(3): e13041. doi:10.1111/psyp.13041

Williams, P., A. Heathcote, K. Nesbitt and A. Eidels (2016). “Post-error recklessness and the hot hand.” Judgment and Decision making 11(2): 174.

Yeung, N. and A. G. Sanfey (2004). “Independent coding of reward magnitude and valence in the human brain.” J Neurosci 24(28): 6258–6264. doi:10.1523/JNEUROSCI.4537-03.2004

Yeung, N. and C. Summerfield (2012). “Metacognition in human decision-making: confidence and error monitoring.” Philos Trans R Soc Lond B Biol Sci 367(1594): 1310–1321. doi:10.1098/rstb.2011.0416

